# Establishment and comparison of three kinds of chronic bacterial cystitis models in mice

**DOI:** 10.1101/2025.07.13.664604

**Authors:** Wan-Bing Chen, Fu Lv, Jing Shao, Zhi Guo, Kun-Yi Wu

## Abstract

Cystitis is a common urinary system disease in women. However, there are currently no effective treatment options except antibiotics. Therefore, establishing an effective and stable animal model is helpful for further study of cystitis. In this study, we established three chronic cystitis models from C57BL/6 female mice inoculated with urinary tract pathogenic Escherichia UTI89 through urethral perfusion: CPP co-infection group, seven days interval repeated infection group, and two consecutive days repeated infection group. The three models were successfully constructed, and the survival rates of the three models were 66%, 75%, 100%, respectively. The bladder infection rates of the surviving mice were 100%, 100%, 50%, respectively. H&E staining showed that the inflammation and damage were the most severe in seven days interval repeated infection group. Sirius red staining showed that the degree of fibrosis was the highest in CPP co-infection group. Immunofluorescence staining of bladder inflammatory cells showed that inflammatory cells and macrophages in the CPP co-infection group and the seven days interval repeated infection group expressed more. Basal layer epithelial cell staining showed that the epithelial cells of the three infection models had different degrees of proliferation. The three strategies can all prepare mouse chronic cystitis models with their own characteristics, and these models can be used for experimental studies related to different levels of bladder infection and fibrosis, laying a foundation for further research on the mechanism of chronic cystitis.

## Introduction

In the era of rapid medical science and technology, infectious diseases are still threatening the life safety and health of people. Urinary tract infections (UTIs) are one of the most common infectious diseases, mainly affecting women and children. According to statistics, about 40-50% of women have at least one urinary tract infection in their lifetime, and 20-30% of women have a recurrence 3-4 months after the initial infection[1]. UTIs is mainly caused by urinary tract pathogenic Escherichia, and the incidence of women is significantly higher than men[2]. Cystitis is a common urinary tract infection disease, accounting for about 60% of urinary tract infections, the main characteristics are bladder pain, frequent urination and hematuria[3]. At present, antibiotics are the main treatment for bacterial cystitis, but with the increasing resistance of bacteria, it is urgent to explore more effective treatment strategies. There are few studies on the pathogenesis and treatment of bacterial cystitis, so it is of great significance to establish a reliable animal model for the study of this disease.

In this study, three chronic bacterial cystitis models were constructed by injecting urinary tract pathogenic Escherichia coli UTI89 into urinary, and the pathological changes of bladder tissues in different models were evaluated and compared. As a new research method, these models can be applied to the study of pathogenic mechanism and treatment of chronic bacterial cystitis.

## Materials and methods

### Mice

SPF-grade sterile C57BL/6 female mice aged 10-12 weeks and weighing 20-22 g were used for the experiment, provided by the Animal Laboratory Center of Xi ‘an Jiaotong University School of Medicine, License No. SYXK (Shaanxi) 2015-0002. All animal experiments have been approved by the Animal Experiment Ethics Committee of Xi ‘an Jiaotong University (Approval number: XJTUAE2023-2331).

### Reagents and instruments

Cyclophosphamide (CPP, C7397-1, Sigma-Aldrich, USA), LB liquid medium and Cystine Lactose Electrolyte Deficient (CLED) medium (Oxoid, USA), Hematoxylin (C0107, Beyotime, China), Eosin (C0109, Beyotime, China), Sirius red (BP-DL029,SBJbio,China), Cytokeratin 5 Polyclonal (KRT5) antibody (28506-1-AP,Proteintech,USA),CD45(103120, Biolegend, USA), Ly6G (127620,Biolegend, USA),DAPI (C1002,Beyotime,China), Fluor™594 Donkey Anti-Rat lgG (H+L) (34412ES60,YEASEN,China),Fluor™594 Goat Anti-Rabbit lgG (H+L) (33112ES60,YEASEN,China). Stereo-microscope (XTZ-D, China), Biosafety cabinet (BIOBASE BSC-1100, China), centrifuge (Eppendorf 5810R, Germany), frozen microslicer (Leica CM1900, Germany), paraffin embedding machine (Leica HistoCore Arcadia, Germany) Germany), Paraffin microtome (Leica RM2235, Germany), optical microscope (Zeiss Axiolab, Germany), Laser confocal microscope (Leica TCS SP8, Germany).

### Establishment of three chronic bacterial cystitis models in mice

Urinary tract pathogenic E. coli UTI89 was presented by Professor Wu Ding Zhou of King’s College London, UK. A mouse model of chronic cystitis was established by injecting UTI89 (50μL with 1×10^8^ CFU/mL bacterial solution in sterile PBS) in the urinary tract of C57BL/6 female mice, with sterile PBS as the control mice[4]. Construction model of three kinds of chronic infective cystitis models in mice (Figure 1). 1) CPP co-infection group (CCI): Mice were intraperitoneally injected with CPP (300 mg/kg) one day before UTI89 inoculation, urinary tract infection model was established the next day, and mice were killed under anesthesia and bladder tissues were collected on the 40th day. 2) 7 days interval repeated infection group (IRI): UTI89 was inoculated into the urethra of mice via urinary tract, and repeated infection was carried out five times, each time at an interval of 7 days. The mice were killed by anesthesia on the 35th day and samples were collected. 3) 2 days consecutive repeated infection group (CRI): UTI89 was inoculated into the urethra of mice via urinary tract, repeated infection was carried out at 24h with the same dose, and the mice were killed by anesthesia on the 28 days and samples were collected.

**Fig 1.**
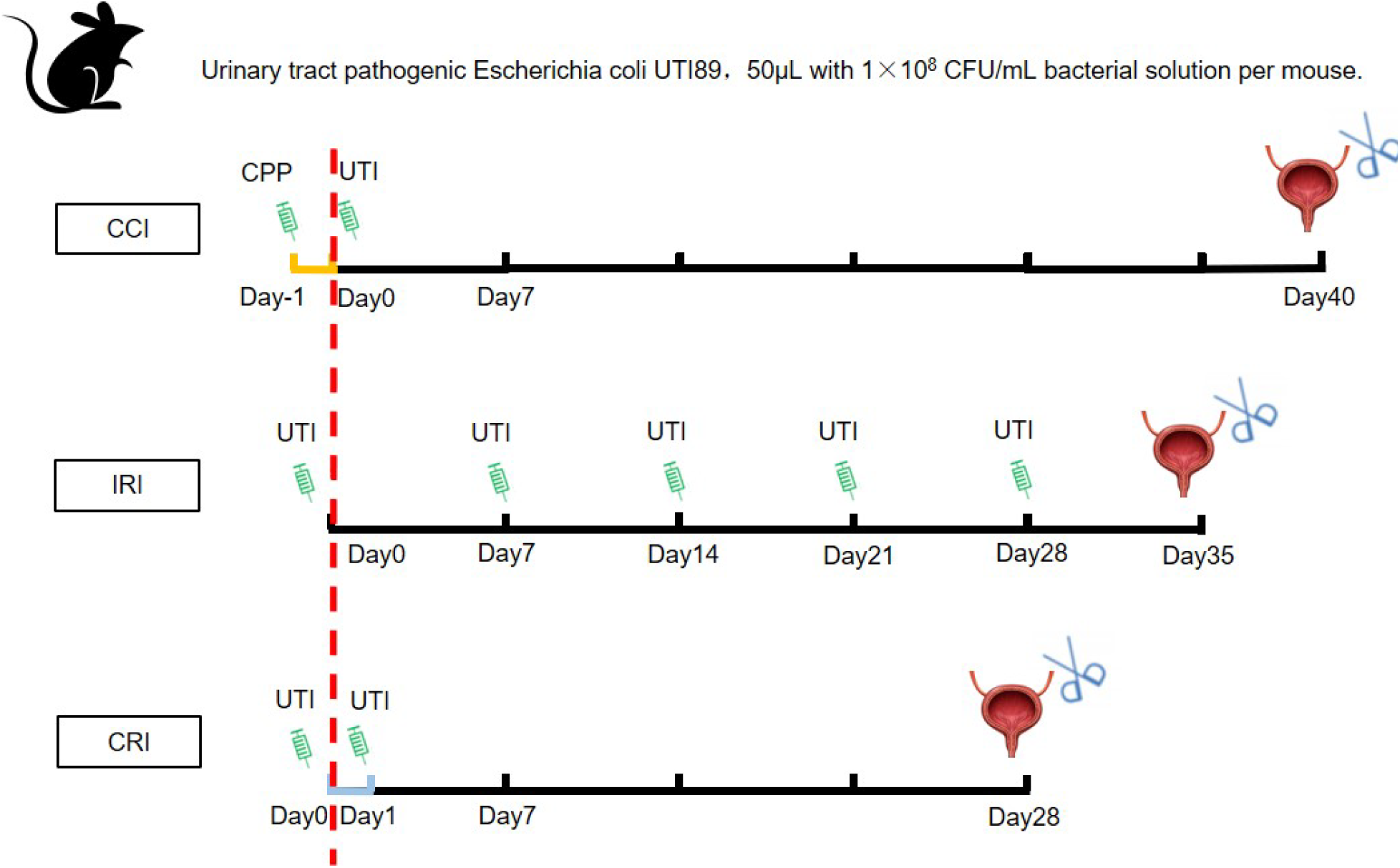
Three models of chronic bacterial cystitis

### Urine colony forming units test (CFU)

Urine colony forming units was detected by bacterial coating method. Mouse urine was collected at different time points, and the urine stock solution was diluted step by step according to 1:10, 1:100 and 1:1000. That is, 10μL urine stock solution was diluted 10 times by adding 90μL sterile PBS, and 10μL urine diluent solution was diluted 100 times by adding 90μL sterile PBS. Then 10μL this of diluent was added to 90μL of sterile PBS and diluted 1000 times. The 50μL diluent at all levels was evenly coated on a crowded plate and incubated at 37°C overnight to count the number of colonies.

### H&E staining

Mouse bladder tissues were collected, fixed overnight with 4% paraformaldehyde, paraffin embedded and sliced (4µm). H&E staining was used to evaluate the histological changes. Paraffin sections were dewaxed with xylene, dehydrated with gradient alcohol, stained with hematoxylin for 7-8 min, differentiated by 1% hydrochloric alcohol/reverted blue with aqueous ammonia, stained with eosin for 15 s, dehydrated and sealed. The bladder tissue score is mainly determined by whether the tissue is edema, whether the epithelium is intact, the degree of thickening of the epithelium, bleeding and the degree of inflammatory cell infiltration. 0: no lesions, no tissue edema, no bleeding, no inflammatory cell infiltration; 1 divided into mild lesions, slight inflammatory cell infiltration, basically no bleeding; 2. A small number of lesions, some inflammatory cell infiltration, a small amount of bleeding, edema, a small amount of epithelium thickening; 3: divided into moderate lesions, obvious inflammatory cell infiltration, epithelial cell shedding; 4: large number of lesions, obvious inflammatory cell infiltration, bladder epithelium thickening, obvious bleeding; 5: Severe lesions, inflammatory cells infiltrate the whole tissue, massive bleeding, obvious thickening of the epithelium, epithelial cells shed. The degree of bladder pathological injury was assessed by two pathologists using double-blind method, and was observed and photographed under optical microscope.

### Sirius red staining

Sirius red staining was used to evaluate tissue fibrosis. Paraffin sections were dewaxed with xylene, dehydrated with gradient alcohol, soaked with Sirius red solution for 1 h, stained with hematoxylin for 7-8 min, dehydrated and sealed, observed and photographed under optical microscope, and the collagen deposition area was analyzed using Image J software.

### Immunofluorescence staining

The bladder tissue was frozen embedded, frozen sections were taken for 4μm, fixed with propyl alcohol for 15 min, and rinsed with PBS for 5 min×3 times. The 10% goat or donkey serum was sealed, incubated with CD45(macrophage marker), KRT5 (basal epithelial cell marker) or Ly6G (neutrophil marker) at 4°C overnight, and rinsed with PBS for 5 min×3 times. Then the secondary antibodies were added and incubated at room temperature for 1 h, and rinsed with PBS for 5 min×3 times. The nuclei were labeled with DAPI and rinsed with PBS for 5 min×3 times. The anti-fluorescence quencher plate was photographed by German Leica SP8 laser confocal microscope, and the positive area was analyzed by Image J software.

### Statistics

Each experiment was repeated at least 3 times, data values were expressed as mean ± standard deviation, area score was analyzed by Image J software, experimental data were statistically analyzed by GraphPad Prism 10 software, and differences between groups were analyzed by One-way ANOVA test. P < 0.05 was considered statistically significant.

## Results

### Comparison of three models’ urine CFU

The colony forming units in the urine of mice is an important criterion to evaluate the success of the chronic bacterial cystitis model. When the bacterial load in the urine reaches 1×10^4^ CFU/mL, the cystitis model is considered to be successfully established[5]. After modeling, urine was collected and coated at different time points to calculate the CFU in urine. In the CCI, 1/3 of the mice died during the infection (4 mice survived / 6 mice in total), and the remaining 2/3 of the mice were able to collect urine when the model was collected on the 40th day. In the IRI, 1/4 of the mice died during infection (9 mice survived / 12 mice in total), and the remaining three quarters were able to collect urine when the model was collected on the 35th day. In the CRI, no mice died during the entire infection process (8 mice survived / 8 mice in total) and urine was collected at the 28th day (Figure 2, A). Urine of infected mice was collected and coated. The CFU in urine of mice in CCI and IRI were higher than 1×10^4^ CFU/mL, and the bladder infection incidence rates of the surviving mice were 100%. Among, the urine bacterial content of CCI was 1.163×10^7^ CFU/mL, 1.123×10^7^ CFU/mL, 1.07×10^7^ CFU/mL, and 1.187×10^6^ CFU/mL, respectively. The bacterial content of urine in the IRI was 1.35×10^7^ CFU/mL, 4.5×10^4^ CFU/mL, 1.45×10^4^ CFU/mL, 4×10^4^ CFU/mL, 3.67×10^4^ CFU/mL, 7.89×10^6^ CFU/mL, 4.53×10^4^ CFU/mL, 6.57×10^4^ CFU/mL, 2.15×10^5^ CFU/mL, respectively. But in the CRI, only 1/2 of the mice had urine bacterial content higher than 1×10^4^ CFU/mL, and the bladder infection incidence rates of the surviving mice was 50%. The urine bacterial content at the time of collection was 9.82×10^5^ CFU/mL, 1.15×10^4^ CFU/mL, and 2.68×10^4^ CFU/mL, 3×10^4^ CFU/mL respectively, and no bacteria were detected in the remaining four mice (Figure 2, B and C). The results showed that although the mice in the CCI and IRI died during the infection process, the surviving mice were able to form models, while the mice in the CRI did not die during the infection process, but only half of the models were formed.

**Fig 2.**
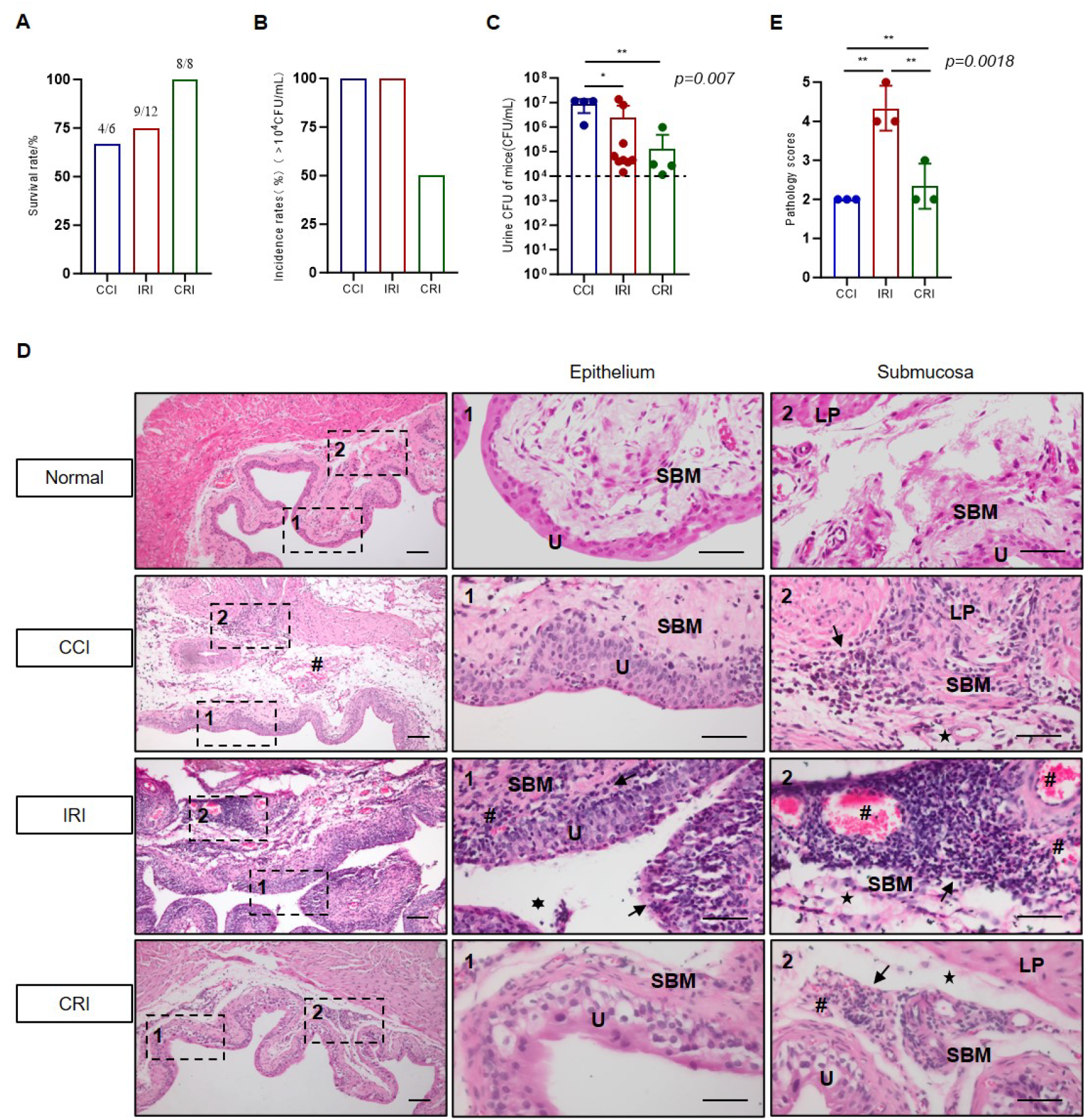
Comparison of urine bacterial load and pathology in three chronic bacterial cystitis models. (A) The survival rate of mice in the three models (survival rate = number of mice alive at the time of collection/number of mice at the time of modeling ×100%), the number of mice in each group was 6, 12 and 8, respectively. (B) The bladder infection incidence rates of the surviving mice with chronic bacterial cystitis (the number of mice receiving urine bacteria greater than 1×104 CFU/mL/the number of mice receiving urine ×100%) was 4, 9 and 8 mice in each group, respectively. (C) Urine CFU of mice with chronic bacterial cystitis. (D) H&E staining representative images of bladder sections in the normal group and the three cystitis models group. Left: Bladder tissue section taken at 10x, scale: 100μm; Middle and right: Bladder tissue section taken at 40x, scale: 50μm. Symbol: Five-pointed star: tissue edema; Hexagonal star: epithelial exfoliation; #: bleeding; Arrow: infiltrating inflammatory cells; U: bladder epithelium; SBM: submucosa; LP: Lamina propria. (E) Pathology scores of three cystitis models. The score was based on tissue edema, epithelial integrity, epithelial thickening, bleeding, and inflammatory cell infiltration. 0, no lesions; 1, mild lesions; 2, a small number of lesions; 3, moderate lesions; 4, a large number of lesions; 5. Severe lesions. The data were analyzed by One-way ANOVA test. *, P<0.05; **, P<0.005. n=3 mice.

### Comparison of three models’ bladder injury degree

In order to explore the pathological changes of chronic bacterial cystitis, the bladder tissues of three mice in each group of cystitis were randomly selected for H&E staining (Figure 2, D). The results showed that the bladder wall and bladder epithelium of normal mice bladder tissue were intact, without obvious inflammatory cell infiltration, vacuole, tissue edema and obvious bleeding. In the CCI, thickening of bladder epithelium, infiltration of some inflammatory cells, tissue edema and bleeding were observed. In the IRI, a large number of inflammatory cells were infiltrated, the bladder epithelium was obviously shed, the bladder epithelium thickened, the submucosa bleeding was more, and tissue edema was observed. In the CRI, there was a small amount of inflammatory cell infiltration, mild tissue edema, a small amount of bleeding in the submucosa, and a small amount of thickening of the bladder epithelium. Through the pathological score of the three models, it was found that the pathological injury was the most severe in the IRI (Figure 2, E).

### Comparison of three models’ bladder tissue inflammatory cell infiltration

In order to further compare the inflammatory conditions of the three models, we performed fluorescent staining of CD45 inflammatory cell immunity in the bladder tissue of the three models. CD45 positive cells (red) were mainly distributed in the upper cortex and submucosa in the CCI, and the proportion of CD45+ staining area was 4.450% (Figure 3, A). There were more CD45 positive cells in the IRI, indicating a more severe inflammatory response, which was mainly distributed in the submucosa and distributed in clusters, and the proportion of CD45 positive cells staining area was 5.730% (Figure 4, B). There were significantly fewer CD45 positive cells in the CRI, and the proportion of CD45 positive cells staining area was 0.809% (Figure 3, C). The statistical results of CD45 positive cells showed that the leukocyte infiltration was more serious in the IRI (Figure 3, D).

**Fig 3.**
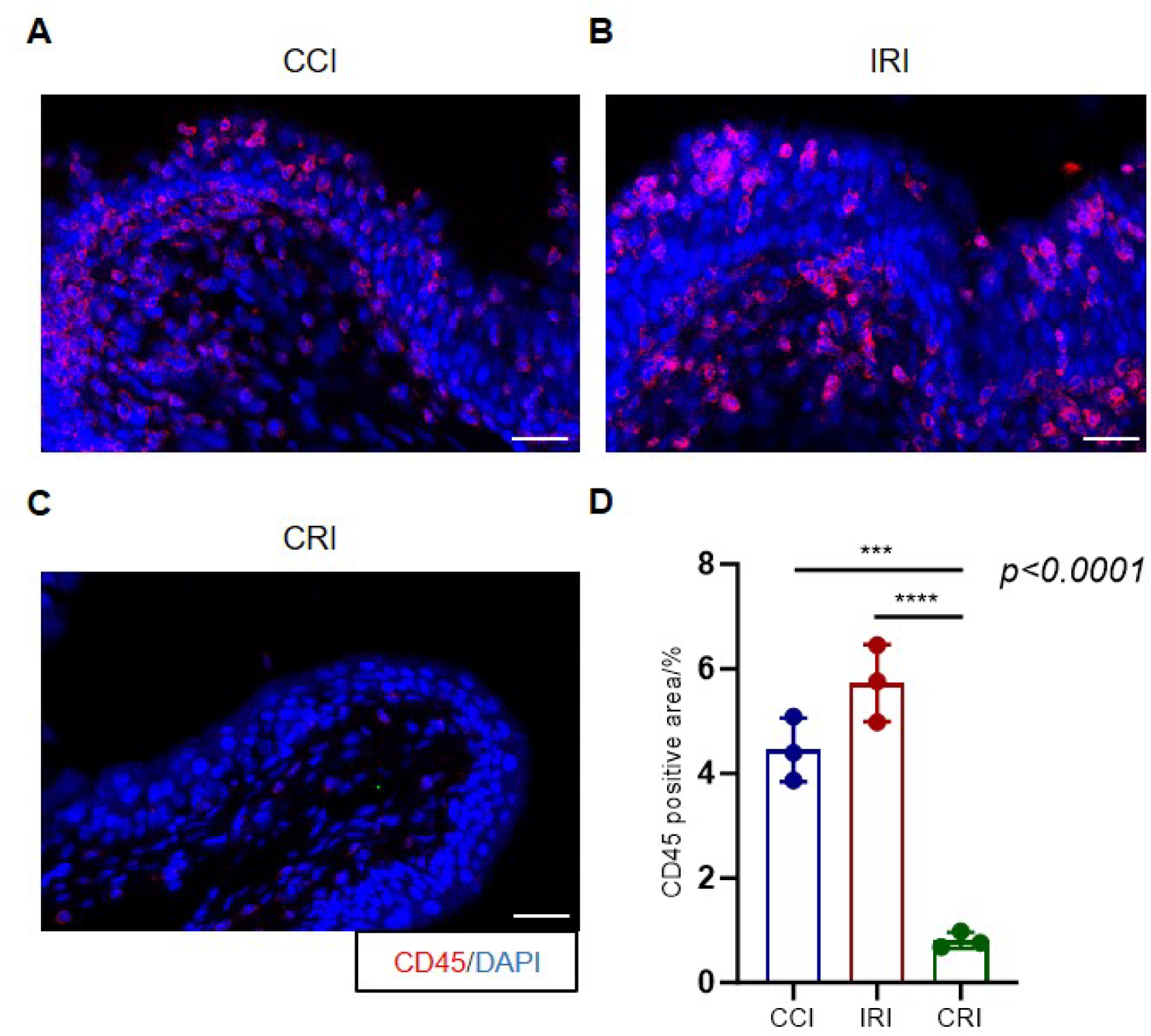
Macrophage immunofluorescence staining in three chronic bacterial cystitis models. (A) CCI. (B) IRI. (C) CRI. Red: CD45 macrophage staining. Blue: DAPI nuclear staining. Scale bar: 25μm. (D) CD45 staining positive area percentage. The data were analyzed by One-way ANOVA test. ***, P<0.001; ****, P<0.0001. n=3 mice.

**Fig 4.**
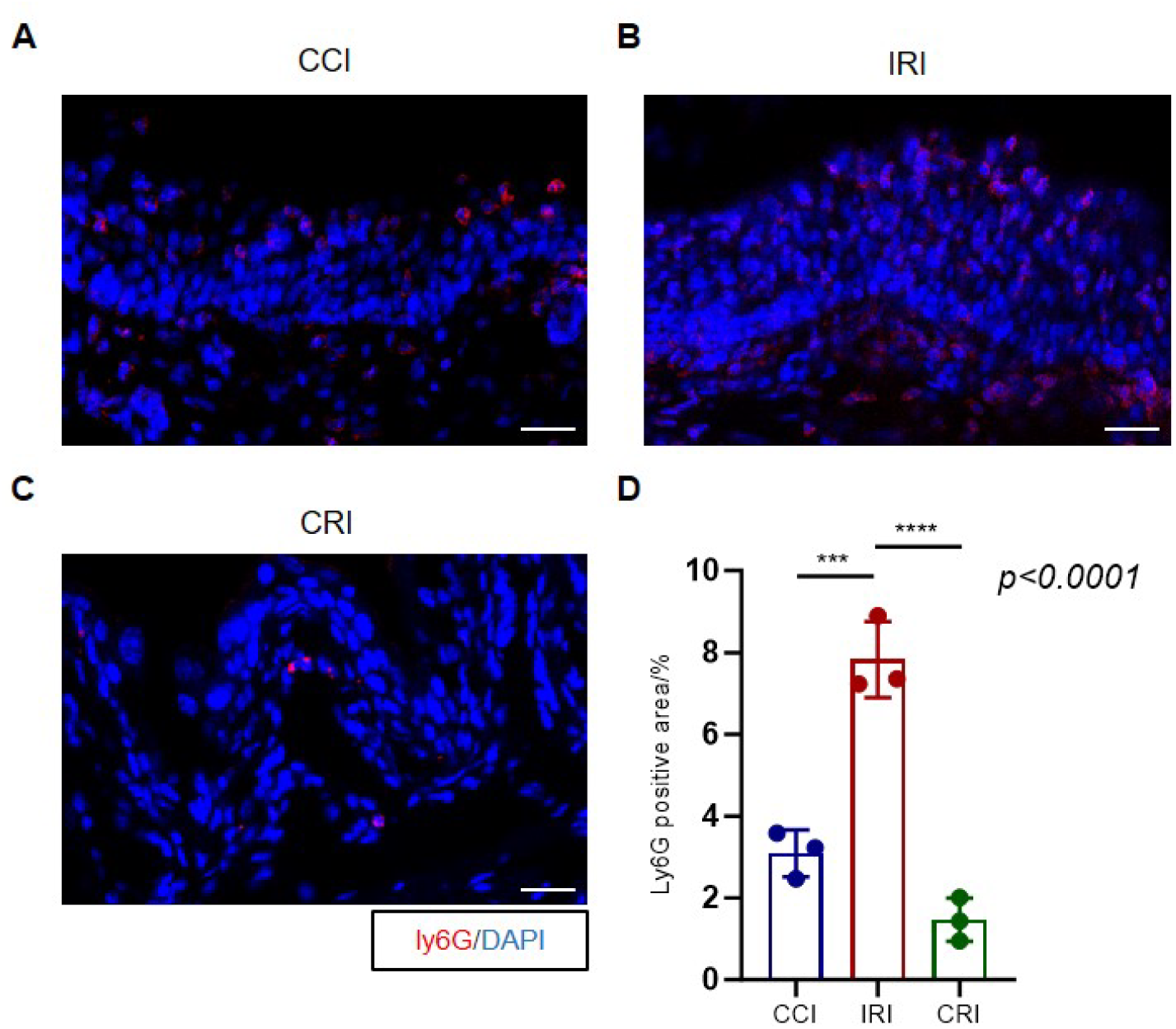
Neutrophil immunofluorescence staining in three chronic bacterial cystitis models. (A) CCI. (B) IRI. (C) CRI. Red: Ly6G neutrophil staining. Blue: DAPI nuclear staining. Scale bar: 25μm. (D) Ly6G staining positive area percentage. The data were analyzed by One-way ANOVA test. ***, P<0.001; ****, P<0.0001. n=3 mice.

Bladder tissue neutrophils are one of the most abundant immune cells in the bladder, and studies have shown that neutrophils play a key role in infection, immunotherapy, and inflammatory responses. To further compare the inflammation of the three models, Ly6G neutrophil fluorescence staining (red) was performed on the bladder tissues of the three models respectively. The results showed that Ly6G positive cells were mainly distributed in the bladder epithelium and less in the submucosa in the CCI, and the average proportion of Ly6G positive staining area was 3.09% (Figure 4, A). In the IRI, there were more Ly6G positive cells, which were mainly distributed in the bladder epithelium and submucosa and expressed in clusters. The average proportion of Ly6G positive cells staining area was 7.83%, suggesting that the inflammatory response was more severe in this group (Figure 4, B). In the CRI, there were fewer Ly6G positive cells, and the average proportion of Ly6G positive cells staining area was 1.47% (Figure 4, C). The statistical results of Ly6G positive cells showed that the cell infiltration was more serious in the IRI, followed by the CCI, and the CRI was the lightest (Figure 4, D).

### Comparison of three models’ bladder tissue collagen deposition

The extracellular matrix and fibrosis of bladder tissue were evaluated by Sirius red staining. The collagen deposition of the three cystitis models were mainly distributed in the submucosa and muscle layer, and the degree of fibrosis was as follows: CCI>IRI>CRI (Figure 5, A-D). The statistical analysis of the positive area of Sirius red staining showed that normal tissues was the least (the average area was 7.16%), the CCI was 60.92%, the IRI was 44.14%, and the CRI was 20.19% (Figure 5, E). The results indicate that chronic bacterial infection can lead to cystic fibrosis, and CPP as an immunosuppressive agent may promote the occurrence and development of cystic fibrosis.

**Fig 5.**
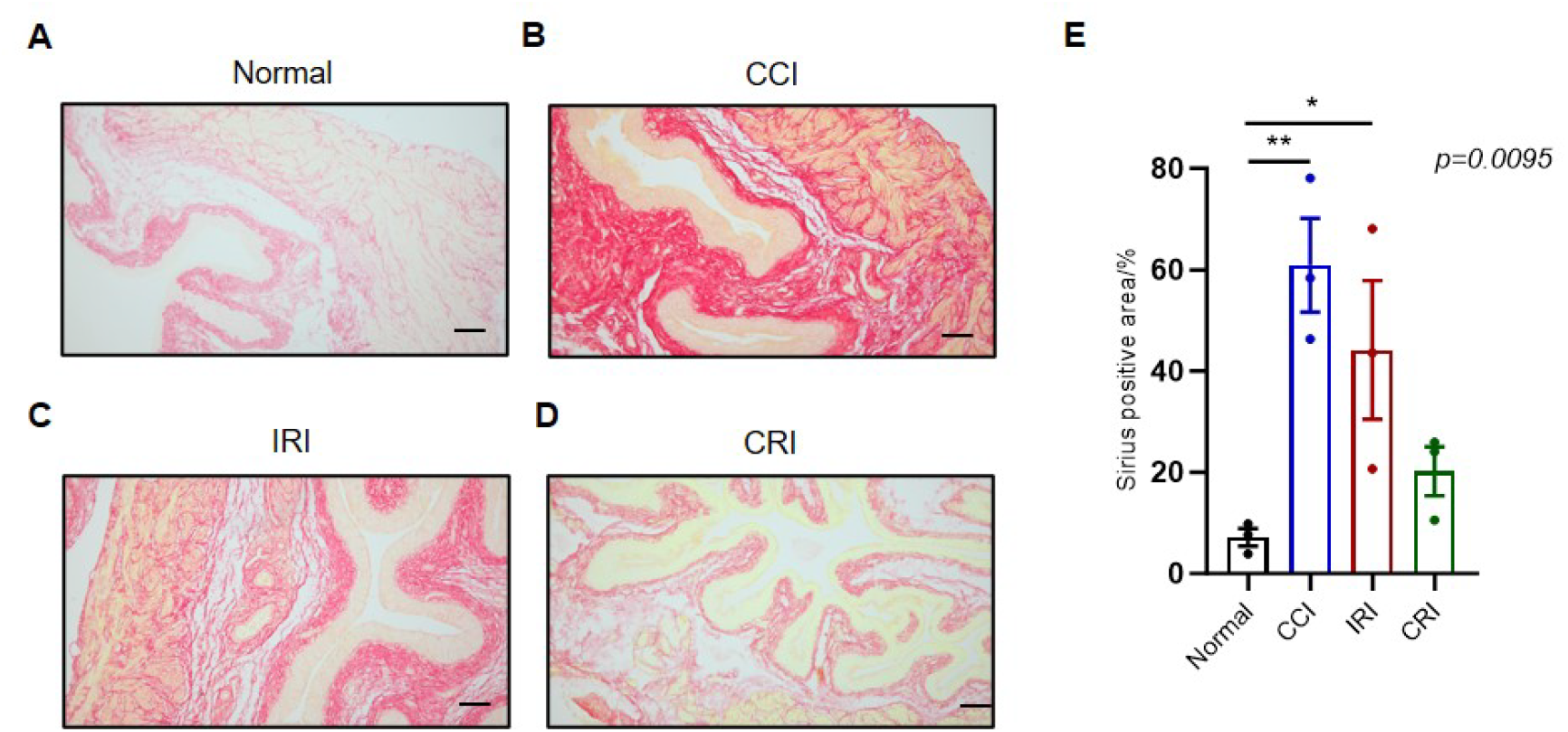
Sirius red staining of three chronic bacterial cystitis models. (A) Normal group. (B) CCI. (C) IRI. (D) CRI. Bladder tissue section taken at 10x. Scale bar: 100 μm. (E) Sirius-red positive staining area of normal tissue and three cystitis model bladder sections. The data were analyzed by One-way ANOVA test. *, P<0.05; **, P<0.005. n=3 mice.

### Comparison of three models’ bladder epithelial cell proliferation

Epithelial cell proliferation is a typical pathological manifestation of chronic cystitis. In order to determine the epithelial proliferation of the three models, the bladder tissues of the three models were respectively stained with KRT5 (epithelial basal layer cell marker, red) (Figure 6, A-D). Then five distances between the bladder epithelium were randomly selected to measure and the proliferative thickness of epithelial cells was calculated. Compared with normal bladder tissue (mean thickness 31.2μm), all three cystitis models showed significant epithelial thickening. The bladder epithelium thickened the most in the IRI (mean thickness 61.3μm), followed by the CCI (mean thickness 58μm) and the CRI (mean thickness 51.8μm) (Figure 6, E). The results suggested that repeated bacterial infection could lead to the thickening of the bladder epithelium, and the proliferation of epithelial cells appeared in different degrees in all three models.

**Fig 6.**
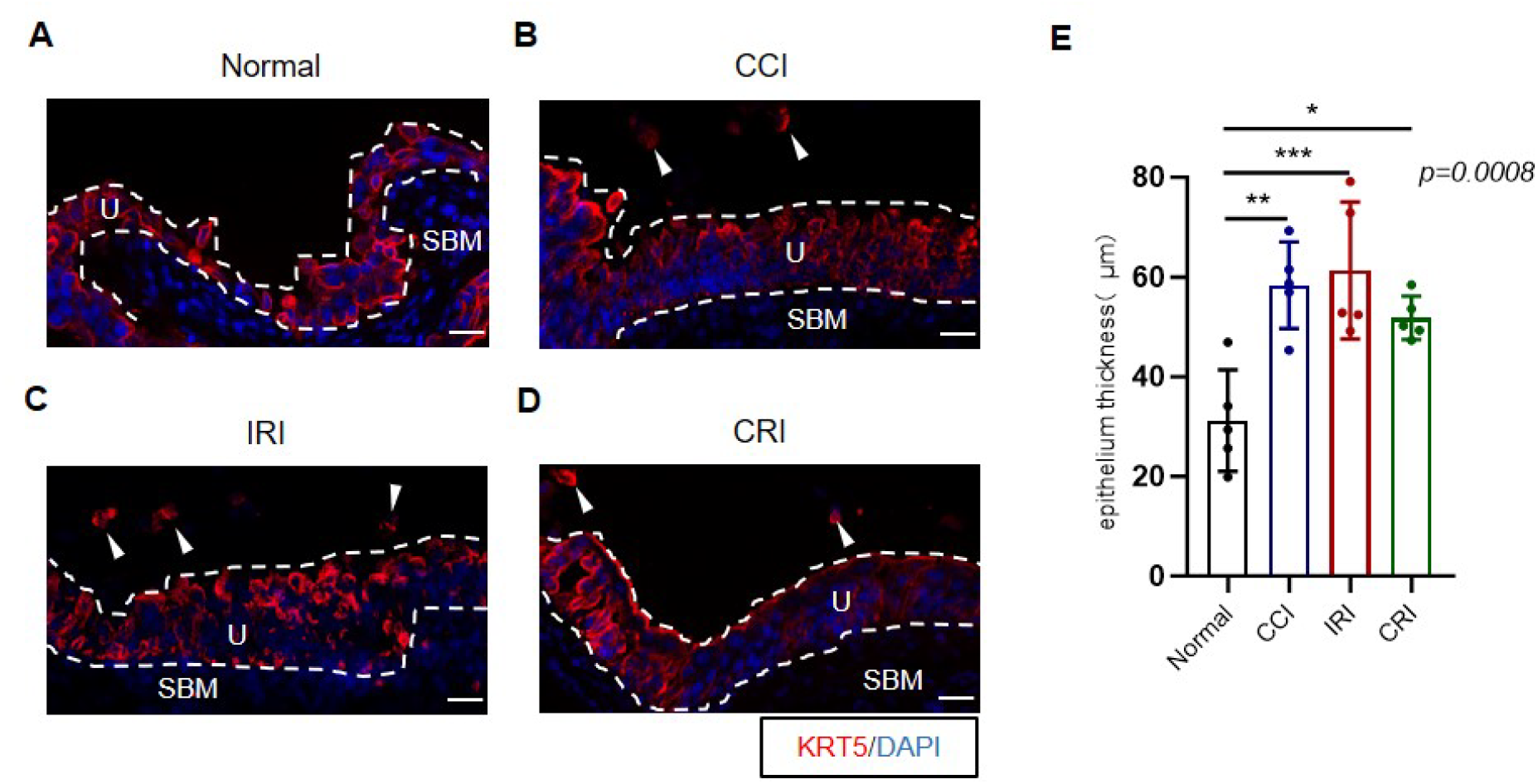
KRT5 fluorescence staining of three chronic bacterial cystitis models. (A) Normal group. (B) CCI. (C) IRI. (D) CRI. Blue: DAPI nuclear staining. U: bladder epithelium; SBM: Submucosa. White arrow: Shed epithelial cells. Scale bar: 25μm. (E) Average bladder epithelium thickness. The data were analyzed by One-way ANOVA test. *, P<0.05; **, P<0.005; ***, P<0.001. n=5 mice.

## Discussion

Urinary tract infections (UTIs) are one of the most common bacterial infectious diseases in the clinic. Most UTIs are caused by UPEC migrating from the intestine to the urethra and eventually to the bladder[6, 7]. UPEC can form bacterial clusters in mouse bladder epithelial cells and lurk in the deep part of bladder tissue, leading to repeated bladder infection[8]. At present, the most common method of bacterial cystitis model is to simulate the pathophysiological process of clinical human urinary tract infection by injecting bacteria into mouse urethra[9, 10]. In this study, three models of chronic bacterial cystitis were established using the modeling method of retrograde urethral infection mastered by our research group[11, 12], and the characteristics of survival rate, incidence rate, pathological injury, inflammatory response, tissue fibrosis and repair degree of epithelial cell injury of the three models of mice were compared, aiming to provide the optimal model scheme for exploring the mechanism of chronic bladder infection.

Chronic bacterial cystitis in mice is characterized by persistent presence of high titer pathogens in the urine, accompanied by histological changes in the bladder epithelium and lamina lamina, such as bladder epithelial hyperplasia with lack of terminal differentiation of surface umbrella cells and submucosal lymph aggregation[13]. This study established and analyzed the survival rate, incidence rate and histomorphology of three kinds of infected mice, and the survival rate was: CRI>IRI> CCI. The incidence rate was: CRI=IRI> CCI. The degree of pathological injury was: IRI> CCI> CRI. The results showed that the three models met the standard of chronic bacterial cystitis, but their pathological manifestations were different. Previous studies have found that bladder stress, urination frequency and inflammatory response are increased when low concentrations of CPP are used in mice, and the mechanism of damage repair can be explored by inducing persistent bladder inflammation through CPP[14]. The results confirmed that CPP combined with infection can simulate chronic cystitis well, but CPP as an immunosuppressant may affect the immune microenvironment in vivo. The IRI and the CRI are mainly simulated clinical recurrent infection models. Compared with the CRI,the IRI has more severe bladder injury and more significant epithelial thickening, which is more suitable for drug therapy research.

Different infection routes and strategies have different effects on the body. The three models used in this study showed different inflammatory responses despite the same amount of inoculated bacteria. Excessive inflammatory cells lead to bladder tissue damage, which allows bacteria to colonize continuously[15]. The results of immunofluorescence staining of macrophage and neutrophil showed that the three models of epithelial cells had different degrees of inflammation. Compared with normal bladder tissue, the infiltration degree of bladder inflammatory cells was the highest in the IRI, followed by the CCI, and the inflammation degree was relatively light in the CRI. Long-term and sustained inflammatory response is considered to be one of the main driving forces of fibrosis[16]. The results of Sirius red staining showed that collagen were deposited in the interstitial of the bladder to varying degrees in all three models, indicating the fibrosis. The degree of fibrosis was relatively light in the CRI, and the degree of fibrosis was the highest in the CCI, which may be due to the effect of CPP on promoting tissue fibrosis[17, 18]. External injury such as infection can activate the proliferation, differentiation and repair function of bladder urothelial cells[19]. KRT5 is a marker of basal epithelial cells, which can reflect the proliferation of epithelial cells. Compared with the normal bladder, the epithelial cells of the three models were thickened to varying degrees, and the proliferation of epithelial cells increased significantly in the IRI, while the abnormal proliferation of bladder epithelial cells would lead to the reduction of urine storage in the bladder and thus frequent urination[20]. This study also has some limitations, such as the limited number of mice in the three models, and the sample size needs to be expanded in the future to further confirm the above conclusions and determine their accuracy and universality.

In summary, in this study, three different chronic bladder infection models were established using the clinical isolation of E. coli UTI89 for retrograde transurethral bladder infection. The characteristics of the models were compared, and the inflammation and fibrosis degree of the three mouse chronic cystitis models were comprehensively evaluated, providing theoretical basis for animal experiments to further study the mechanism and pharmacodynamic evaluation of chronic bacterial cystitis.

## Acknowledgements

Wan-Bing Chen: Conduct research, draft articles, statistical analysis; Jing Shao: Experiment implementation and data collection; Fu Lu: Data collection; Zhi Guo: Experiment implementation; Kun-Yi Wu: Design experiment, paper revision, research guidance, financial support.

## Data sharing statement

All data are available in the main text or the Supplementary Material. The custom resources related to the manuscript are available from the corresponding authors upon request. This article does not report large datasets, original code, and reanalysed data.

